# A developmental pathway for epithelial-to-motoneuron transformation in *C. elegans*

**DOI:** 10.1101/2022.05.27.493712

**Authors:** Alina Rashid, Maya Tevlin, Yun Lu, Shai Shaham

## Abstract

Motoneurons and motoneuron-like pancreatic beta cells arise from radial glia and ductal cells, both tube-lining progenitors that share molecular regulators. To uncover programs underlying motoneuron formation, we studied a similar, cell-division-independent transformation of the *C. elegans* tube-lining Y cell into the PDA motoneuron. We find that *lin-12/Notch* acts through *ngn-1/Ngn* and its regulator *hlh-16/Olig* to control transformation timing. *lin-12* loss blocks transformation, while *lin-12*(gf) promotes precocious PDA formation. Early basal expression of both *ngn-1/Ngn* and *hlh-16/Olig* depends on *sem-4/Sall* and *egl-5/Hox*. Later, coincident with Y-cell morphological changes, *ngn-1/Ngn* expression is upregulated in a *sem-4/Sall* and *egl-5/Hox*-dependent but hlh-16/Olig-independent manner. Subsequently, Y-cell retrograde extension forms an anchored process priming PDA axon extension. Extension requires ngn-1-dependent expression of the cytoskeleton organizers UNC-119, UNC-44/ANK, and UNC-33/CRMP, which also, unexpectedly, activate PDA terminal-gene expression. Our findings reveal key cell-division-independent regulatory events leading to motoneuron generation, suggesting a conserved pathway for epithelial-to-motoneuron/motoneuron-like differentiation.

## INTRODUCTION

During animal development, neurons and neuron-like cells are generated from progenitor cells that often line tubes. For example, in the developing spinal cord, basal processes of radial glial stem cells line the fluid-filled spinal canal. NOTCH/DELTA signaling is implicated in the differentiation of these cells into motoneurons through control of their proliferative state (de la Pompa et al., 1997; Appel et al., 2001; Tan et al., 2016). Down-regulation of the NOTCH-dependent HES1 and HES5 transcriptional inhibitors in differentiating cells induces expression of the proneural bHLH transcription factors OLIG2 and NGN2 (Sagner et al., 2018; Hatakeyama et al., 2004; Ishibashi et al., 1995; Ohtsuka, 1999), leading to adherens junction loss, delamination from the epithelium, cell migration, and neuronal maturation (Ma et al., 1998; Fode et al., 1998; Mizuguchi et al., 2001; Lee et al., 2005). A similar sequence of events characterizes formation of pancreatic insulin-secreting β-cells, which are innervated (Nicolls, 2004) and express genes also active in motoneurons (Thor et al., 1991; Ericson et al., 1992; Polak et al., 1993; Cirulli et al., 1994; Naya et al., 1995; Sommer et al., 1996; Sussel et al., 1998; Gradwohl et al., 2000; Muñoz-Bravo et al., 2013). Here, epithelial progenitor cells lining pancreatic ducts express high levels of NOTCH, which activates HES1 repressor, in turn blocking NGN3 expression (Apelqvist et al., 1999; Jensen et al., 2000; Murtaugh et al., 2003; Shih et al., 2012; Ninov et al., 2012). Down-regulation of NOTCH drives delamination, cell migration, and subsequent differentiation by lifting inhibition of NGN3 (Zhou et al., 2007; Rubio-Cabezas et al., 2011; Gradwohl et al., 2000; Bankaitis et al., 2018), and by allowing NOTCH to directly activate NGN3 expression (Cras-Méneur et al., 2009; Shih et al., 2012). The OLIG family bHLH factor bHLHb4 is expressed in delaminating/migrating cells, and may be involved in β-cell differentiation (Bramblett et al., 2002).

The dynamics of gene expression and the regulatory interactions among genes driving motoneuron formation are not fully understood. Furthermore, distinguishing whether genes governing this differentiation event control cell division or cell fate acquisition can be challenging. To address these issues, we have studied a similar transformation process in the genetically amenable nematode, *Caenorhabditis elegans*.

The *C. elegans* Y cell is one of six epithelial cells that line the rectal tube in first-larval-stage (L1) animals (Figure 1A). In L2 animals, Y loses apical junctions with neighboring cells, migrates anterodorsally, and transforms into the PDA motor neuron (Sulston et al., 1983; Jarriault et al., 2008), which innervates the intestinal and anal depressor muscles (White et al., 1986). Remarkably, this transformation takes place without cell division (Sulston et al., 1983). Forward and reverse genetic studies identified a number of genes required for the Y-to-PDA transition, including *lin-12/Notch, sem-4/Sall, egl-5/Hox*, and several chromatin remodeling genes (Jarriault et al., 2008; Richard et al., 2011; Kagias et al., 2012; Zuryn et al., 2014). Thus, the Y-to-PDA transition is an excellent setting in which to identify regulators that specifically affect motoneuron progenitor-cell differentiation, and not cell division.

**Figure 1:**
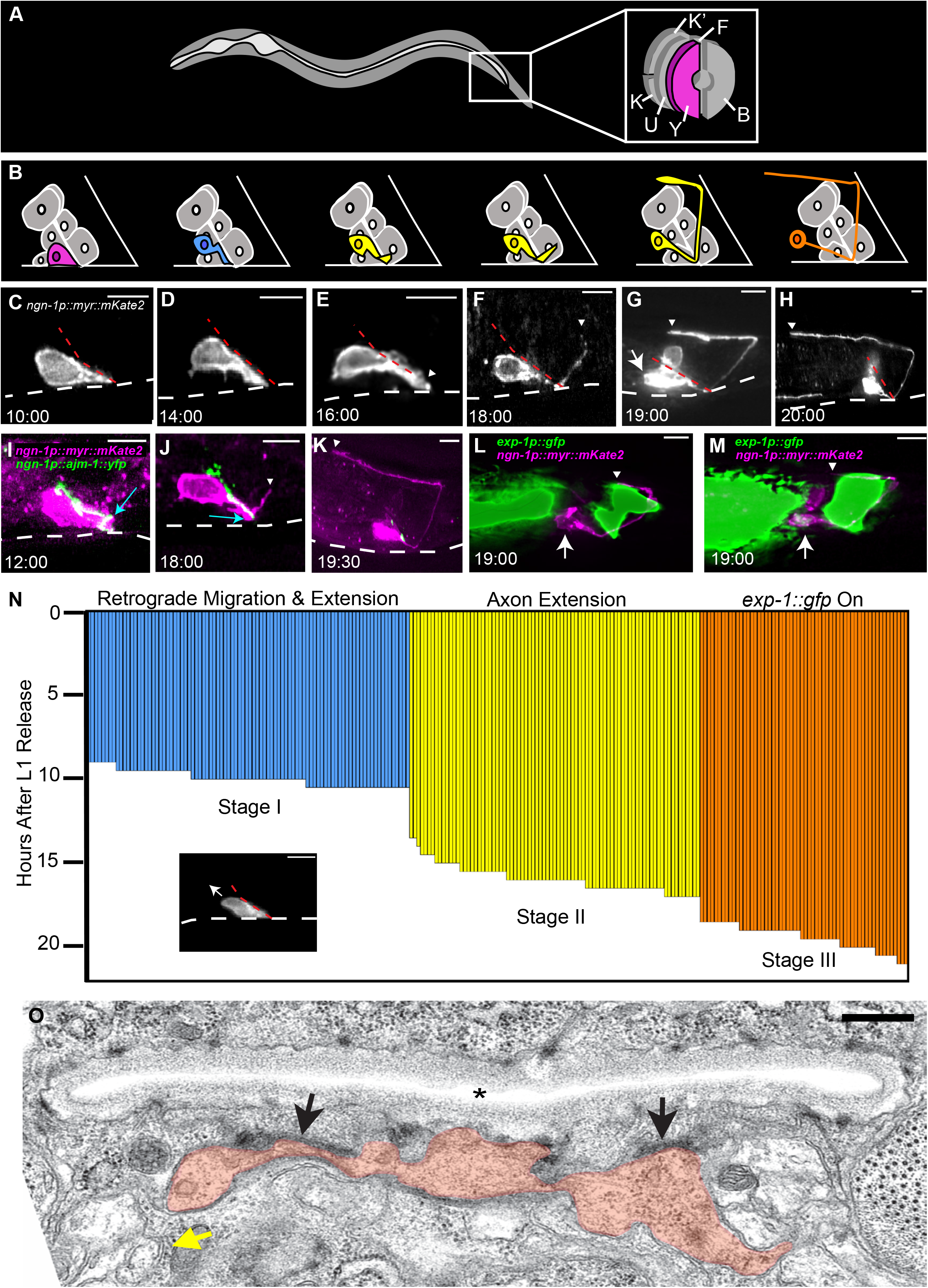
Three morphological stages characterize Y-to-PDA transformation. See also Figures S1, S2, and Table S2. (A) Cellular anatomy of rectal tube In an L1 animal. The Y cell is at the opening. (B) Schematic of Y-to-PDA transformation time course shown in C-H. Y/PDA is pseudo colored to match panel N. (C-M) Time stamps, hours after release from L1 arrest. (C) The Y cell before transformation begins. (D) Anterodorsal cell body movement with cell tip anchored at rectal slit. (E) Axon outgrowth initiation. (F-H) Stages of axon extension. (I-K) Apical junctions along the cell length marked by AJM-1::YFP (green) expressed in Y cell (magenta) are maintained during axon outgrowth (I,J) and disappear after (K). Apical junctions at the rectal slit base (blue arrows) are initially present in the Y cell (I) but disappear during axon outgrowth (J). (L,M) *exp-1p::gfp* expression onset occurs after axon outgrowth. (C-M) Scale bars, 5 μm. Dashed white lines, ventral surface of animal. Dashed red lines, rectal slit. Arrows, cell body. Arrowheads, axon tip. (N) Each horizontal bar represents Y-to-PDA stage of a single animal. Image shows a representative animal at the onset of retrograde migration and extension stage (Arrow, direction of cell migration). Stage I: n = 89, mean = 9.95 ± 0.47 hrs, Stage II: n = 81, mean = 15.93 ± .8 hrs, Stage III: n = 58, mean = 19.43 ± 0.73 hrs. (O) Electron micrograph of an animal staged as in panel F. Orange, Y/PDA. Asterisk, rectal slit. Black arrows, putative adhesion junctions. Yellow arrow, neural bundle into which the PDA process enters. Scale bar, 500 nm.

Here, we report our identification of novel regulators of Y-to-PDA transformation and explore the interactions among them and previously identified genes. We find that loss of *lin-12/Notch* blocks PDA neuron formation, and gain of *lin-12/Notch* function generates a precocious PDA neuron. Thus, *lin-12/Notch* acts as a timing rheostat for motoneuron generation in a cell-division-independent capacity. *lin-12/Notch* functions, at least in part, by regulating *ngn-1/Ngn*, through control of the bHLH gene *hlh-16/Olig. ngn-1/Ngn* basal expression levels are set early on by *sem-4/Sall, egl-5/Hox*, and *hlh-16/Olig*. Coincident with the onset of Y-cell morphological changes and migration, we find an increase in *ngn-1/Ngn* gene expression, governed by *sem-4/Sall* and *egl-5/Hox*, but not *hlh-16/Olig*. Y-cell migration is accompanied by retrograde extension of a process that remains anchored at the rectal slit. The tip of this process serves as a growth point for the PDA axon. Following axonal growth, we identify the axonal cytoskeleton genes *unc-119, unc-44/Ank*, and *unc-33/Crmp* as targets of *hlh-16/Olig* and *ngn-1/Ngn*, and demonstrate a previously unappreciated role for these genes in regulating the expression of motoneuron terminal differentiation genes.

Our results highlight intriguing similarities with spinal-cord motoneuron and pancreatic islet formation, suggesting a conserved module for the differentiation of tube-lining cells into motoneuron/motoneuron-like progeny.

## RESULTS

### Y-to-PDA transformation occurs in distinct stages through a transient epithelial-neuronal hybrid intermediate

To image the sequence of events leading to PDA formation, we developed a fluorescent reporter expressed in both Y and PDA, as this had not been previously published. We leveraged our genetic studies, identifying *ngn-1* as a relevant gene in Y-to-PDA transformation (see below), to generate an *ngn-1 promoter::myr::mKate2* transgene, containing ~4kb of *ngn-1* upstream regulatory sequences, a myristoylation (*myr*) sequence for membrane localization, and the *mKate2* red fluorescent reporter gene. This reporter is strongly expressed in the Y cell of L1 animals (Figure 1B-H). We synchronized a pool of animals by hatching embryos in the absence of food, resulting in growth arrest at the equivalent of ~2 hours of postembryonic development. Separate groups of animals from this pool were imaged every 30 minutes between 6-22 hours following food introduction. We found that the Y-to-PDA transition occurs in a stereotypical pattern (Figure 1B-H,N). Between 9-10.5 hours, the Y cell begins to migrate anteriorly and dorsally, while its posterior tip remains attached to the rectal slit, thereby forming a thick posterior process (Figure 1C,D,N). These initial events are reminiscent of retrograde extension of sensory-neuron dendrites and nerve-ring axons (Heiman and Shaham, 2009; Singhal and Shaham, 2017; Fan et al., 2019). Axon outgrowth, initiating from the anchored posterior tip, then begins between 13.5-17 hours (Figure 1E-H,N).

Importantly, during the Y-to-PDA transition, the cell does not fully delaminate from the rectal epithelium, as was previously proposed (Richard et al., 2011). Supporting this conclusion, we find that the apical junction marker AJM-1::YFP, expressed specifically in Y, continues to localize along the length of the cell, but not at the base of the rectal slit, while the cell extends its axon (Figure 1I,J). Furthermore, electron micrographs of Y/PDA with a partially extended axon reveal persistent apical junctions with other rectal epithelial cells (Figure 1O). Once the axon is fully extended, the cell loses AJM-1 localization to the rectal slit (Figure 1K). We also observed that the nucleus of the cell retains a single prominent nucleolus, a feature of epithelial cells, and only adopts a neuron-like speckled nucleolar morphology after axon outgrowth is complete (Figure S1) (Sulston and Horvitz, 1977). Thus, Y-to-PDA transformation proceeds through a transitory state exhibiting both epithelial and neuronal characteristics.

The mature PDA neuron expresses the EXP-1/GABRB3 GABA-gated cation channel (Jarriault et al., 2008; Beg and Jorgensen, 2003). Surprisingly, we found that an *exp-1 promoter::gfp* reporter transgene is only detected starting at 18.5-21 hours post hatching, and once the PDA axon has passed the dorsal aspect of the anal depressor muscle (Figure 1L-N). Thus, *exp-1/Gabrb3* expression is turned on after axon extension has initiated. We believe this conclusion is not confounded by a lag in visualization of GFP, as previous studies demonstrate that GFP is detected within 30-60 minutes of gene-expression initiation (e.g. Maurer et al., 2007).

In certain conditions, starvation can alter Y-to-PDA transformation timings and events (Becker et al., 2019). To confirm that this is not the case in our studies, we repeated the time-lapse experiments in non-starved animals. We found that retrograde migration occurs between 10.5 and 12.5 hours after hatching, axon extension occurs between 16 and 18 hours after hatching, and onset of *exp-1::gfp* expression occurs between 19.5 and 22 hours after hatching (Figure S2). Since starvation synchronization arrests animals at ~2 hours after hatching, our protocol does not affect the timing or the nature of Y-to-PDA transformation events.

Taken together, our observations reveal three distinct stages in the transformation of Y to PDA: (I) anterior and dorsal cell-body migration, coupled with posterior process retrograde extension, (II) axon outgrowth initiation, and (III) *exp-1/Gabrb3* gene expression. These stages are temporally non-overlapping, as cells extending axons always undergo cell migration and retrograde extension (n = 81), and cells expressing *exp-1/Gabrb3* always undergo migration, retrograde extension, and axon outgrowth (n = 58; Figure 1N).

### NGN-1/NGN is required early during Y-to-PDA transformation

To identify genes required for the Y-to-PDA transformation, we performed a forward genetic screen, seeking animals that fail to express the *exp-1p::gfp* reporter transgene in adults. Such mutants could affect transitions between any of the three stages we defined. We identified 16 such mutants (Table S1), and through complementation tests and whole-genome sequencing found that three (*ns188, ns189, ns192*) carry defects in the gene *sem-4. sem-4* encodes a SALL-family transcription factor and was previously shown to be required for the Y-to-PDA transition (Jarriault et al., 2008), demonstrating that our screen can uncover *bona fide* Y-to-PDA transformation mutants.

Animals homozygous for the *ns185* allele isolated in our screen have a strong defect in *exp-1p::gfp* expression (*PDAp::gfp;* Figure 2A,D,O). To determine where PDA formation fails, we examined *ns185* mutants carrying an *egl-5 promoter::gfp* transgene. This reporter is normally detected in the Y cell of L1 animals, but is not expressed in PDA neurons (Figure 2B,C) (Jarriault et al., 2008; Zuryn et al., 2014). We found that in *ns185* mutants, the Y cell forms properly, undergoes anterodorsal cell-body movement, and a thick posterior process begins to stretch out. However, an axon is not formed, nucleolar morphology remains epithelial, and the cell constitutively expresses the *egl-5p::gfp* reporter (Figure 2D-F,O). Furthermore, *ns185* mutant animals retain apical junctions along the length and base of the cell into adulthood (Figure 2J,K). Together, these results suggest that the gene affected in *ns185* animals is required for the transition from stage I to stage II of Y-to-PDA transformation (Figure 1N).

**Figure 2:**
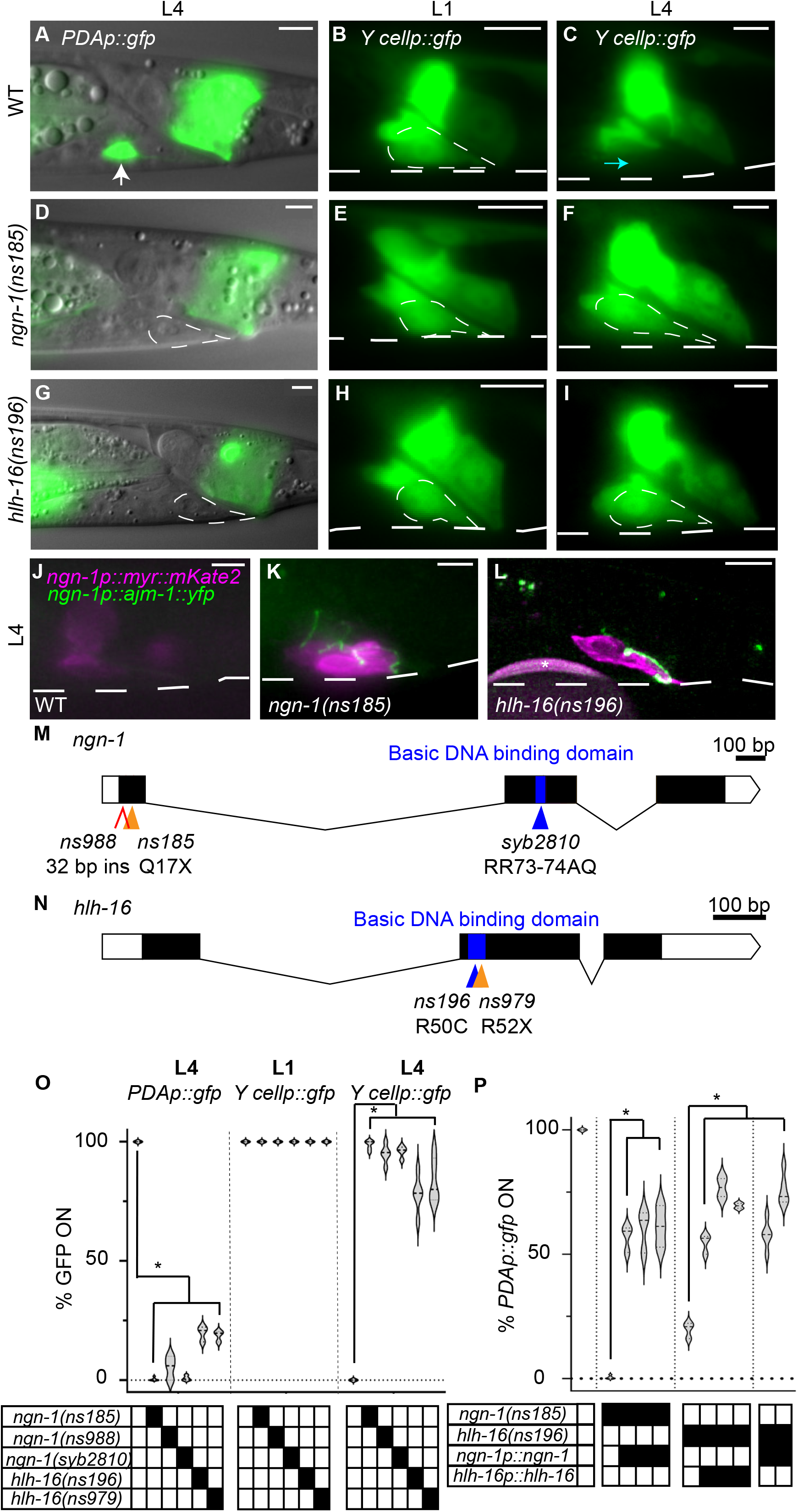
Defective Y-to-PDA transformation in *ngn-1* and *hlh-16* mutants. See also Table S1, S2. (A-C) Wild-type Y-to-PDA transformation. *PDAp::gfp, nsIs131*[*exp-1p::gfp*]. *Y cellp::gfp, bxIs7* [*egl-5p::egl-5(exon1-3)*::*gfp*] reporter. Dashed white circle, outline of Y cell. Scale bars, 5 μm. (A) *PDAp::gfp* expression in L4 PDA (arrow). (B) The Y cell in an L1 animal. (C) Y cell transformed into PDA neuron in L4 animal, *Y cellp::gfp* is off. Cyan arrow indicates where Y cell was in L1 animals. (D-F), same as (A-C), except in *ngn-1(ns185)* mutant. PDA is not formed, and *Y cellp::gfp* is expressed in L4 stage. (G-I), same as (D-F), except in *hlh-16(ns196)* mutant. (J-L) Persistent Y cell in L4 *ngn-1* and *hlh-16* mutants maintain apical junctions. Asterisk, bead used for focusing. (M,N) Amino acid changes are indicated. X, stop codon. (M) The *ns988* allele contains a 32 bp insertion with a stop codon between nucleotides 29 and 30. (O,P) Percent of animals expressing indicated reporters in indicated genotype. Data shown as violin plots. n > 100 animals, n > 3 replicates for all genotypes. Solid black boxes, presence of a mutation or transgene. (O) Significance at p < 0.01, two-way ANOVA with Dunnett’s post-hoc test. (P) Significance at p < 0.01, one-way ANOVA with Dunnett’s post-hoc test.

To identify the affected gene, we used single-nucleotide-polymorphism (SNP) mapping to localize the causal lesion to an 885 kb region on linkage group IV, containing the *ngn-1* gene. A transgene containing ~4 kb of *ngn-1* regulatory sequences, fused to *ngn-1* cDNA, restores PDA neuron formation and *exp-1p::gfp* expression to *ns185* mutants (Figure 2P). Restoration may be incomplete as a result of *ngn-1* overexpression. Sequencing of the gene from *ns185* animals revealed a C-to-T nucleotide change predicted to result in a Q17Stop lesion, which would generate a severely truncated protein (Figure 2M). Furthermore, a CRISPR-generated mutant we isolated, *ns988*, with an early stop codon, phenocopies the defects seen in *ns185* animals (Figures 2M,O). Thus, *ns185* is an allele of *ngn-1*. An additional allele of *ngn-1, ns194*, was isolated from the screen and contains a C-to-T nucleotide change predicted to result in a Q29Stop lesion and a severely truncated protein (Table S1).

*ngn-1* encodes a protein related to vertebrate Neurogenin proteins, a sub-family of basic helix-loop-helix (bHLH) transcriptional regulators that control a variety of developmental processes through DNA binding and helix-loop-helix domain interactions (Sommer et al., 1996; Gradwohl et al., 1996; Ma et al., 1998; Ma et al., 1999; Sun et al., 2001; Grove et al., 2009). To determine whether the DNA-binding domain is required for Y-to-PDA transformation, we used CRISPR-mediated mutagenesis to replace conserved arginine residues (RR) in the NGN-1 basic domain with AQ. This mutation perturbs DNA binding of Neurogenin proteins, while leaving other domains functional (Sun et al., 2001). We found that these *ngn-1(syb2810)* mutants reproduce the defects of *ngn-1(ns185)* and *ngn-1(ns988)* mutants, suggesting that the DNA-binding domain of NGN-1 is required for Y-to-PDA transformation (Figure 2M,O).

To determine where NGN-1 functions, we generated animals carrying a genomically-integrated transgene containing ~4 kb of *ngn-1* regulatory sequences fused to *myr::mKate2* (Figure 1C-H). We found that in the rectal area of early L1 larvae, prior to PDA axon extension, this reporter is expressed exclusively in the Y cell. The same promoter was used to drive *ngn-1* cDNA in our rescue studies (Figure 2P), suggesting that NGN-1 functions cell autonomously in Y for Y-to-PDA transformation.

### HLH-16/OLIG also functions early during Y-to-PDA transformation

*ns196* animals also have a strong defect in *exp-1p::gfp* expression (*PDAp::gfp;* Figure 2G,O). Like *ngn-1(ns185)* mutants, *ns196* animals reliably form the Y cell, which initiates migration and retrograde extension. However, axon formation, nucleolar changes, and silencing of the *egl-5p::gfp* reporter do not take place (Figure 2G-I,O). *ns196* mutant animals also retain their apical junctions along the length and base of the cell (Figure 2L), confirming a failure in the transition from stage I to stage II.

SNP mapping of *ns196* mutants revealed linkage of the causal lesion to a 335 kb region on chromosome I containing the *hlh-16* gene, which encodes a basic helix-loop-helix transcription factor related to vertebrate OLIG2 and bHLHb4 (Reece-Hoyes et al., 2005s; Grove et al., 2009; Bertrand et al., 2011). A transgene containing a 480 bp sequence upstream of the *hlh-16* start codon, fused to the *hlh-16* cDNA and 1.5 kb of *3’* UTR sequences, rescues the mutant defects (Figure 2P). Thus, *ns196* is an allele of *hlh-16*. Indeed, sequencing of the gene from *ns196* mutants revealed a C-to-T nucleotide change leading to a predicted R50C aminoacid substitution in the conserved basic DNA-binding domain (Figure 2N). Thus, the DNA-binding domain of *hlh-16* is required for Y-to-PDA transformation. Furthermore, A CRISPR-generated R52stop mutation *(ns979)* phenocopies the defects seen in *ns196* (Figures 2N,O), confirming *hlh-16* is the relevant gene.

To determine where HLH-16 functions, we first generated animals carrying a genomically-integrated transgene containing 480 bp of *hlh-16* upstream regulatory sequences fused to *myr:gfp. We* did not detect any GFP fluorescence in the rectal area. However, animals carrying a similar transgene, with an additional 1.5 kb fragment of the *hlh-16 3’* UTR, showed specific GFP expression in the Y cell, demonstrating that the *3’* UTR of the gene is important for Y-specific expression (Figure S3F). A construct with the same promoter and UTR driving *hlh-16* cDNA rescues *hlh-16* mutant defects (Figure 2P), suggesting that *hlh-16* functions cell autonomously in Y for the Y-to-PDA transformation.

### *hlh-16/Olig* controls its own expression and basal expression of *ngn-1/Ngn*

As *ngn-1* and *hlh-16* mutants exhibit similar defects, they may function in the same pathway. To address this possibility, we first determined the temporal expression pattern of these genes during the differentiation process. We observed Y-cell fluorescence in animals carrying the *hlh-16p::myr::gfp::hlh-16 3’ UTR* reporter for a 14-hour period following L1 arrest exit, at 2-hour intervals (n > 53 per time point; Figure S4A). Only small changes in expression (~20%) are detected over this time interval. A similar experiment, using the *ngn-1p::myr::mKate2* reporter, also revealed little change in expression over the first 10 hours of imaging. However, from 10-14 hours, as changes in Y-cell morphology become evident, *ngn-1* reporter expression levels nearly double from 73 to 145 a.u. (n > 57 per time point; Figures 3A, S3A). This suggests that *ngn-1* expression is subject to at least two modes of control. Before 10 hours, *ngn-1* is maintained at a static basal level. Between 10-14 hours, expression is boosted. Similar results were obtained in CRISPR-mediated knock-ins into the endogenous *ngn-1* locus, with gfp sequences inserted either directly upstream of the stop codon, or as part of a trans-splicing cassette immediately downstream of the *ngn-1* gene (Figure S3E).

**Figure 3:**
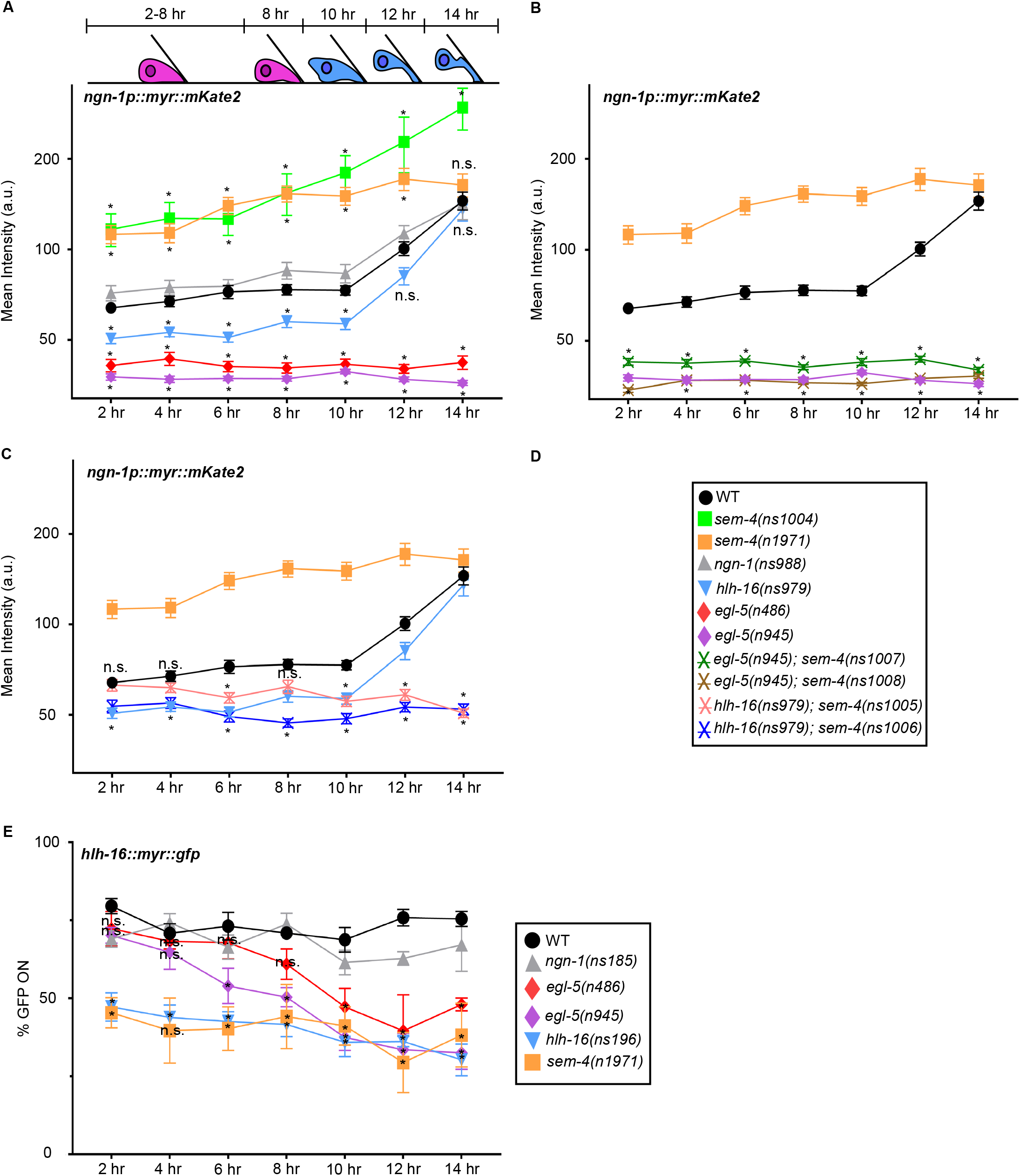
Time course of and effects of mutants on *ngn-1* and *hlh-16* reporter expression. See also Figures S3,S4; Table S2. Schematic depicts Y cell morphology at the indicated time points. Horizontal axis, time after L1 arrest exit. n.s., not significant. (A-C) Y-cell fluorescence levels of *nsIs913* [*ngn-1p::myr::mKate2*] in indicated mutant backgrounds. n > 24 animals per time point. The Y-axis is plotted on a log_2_ scale. (B-C) Significance values only shown for strains not present in panel A. (D) Legend for graphs A-C. (E) As *nsIs943* [*hlh-16p::myr::gfp*] reporter expression is faint, signal quantification is unreliable. We therefore scored the percentage of animals where any fluorescence signal was detected in the Y cell of indicated mutants. Statistics for *ngn-1* mutants are not plotted for clarity. *ngn-1* mutants do not show statistically significant differences from WT animals. n > 5 experiments for all genotypes, n > 15 animals per time point per experiment. Two-way ANOVA with Dunnett’s post-hoc test. Significance at p < 0.05. Error bars, SEM. Statistics in Table S2.

We next investigated *hlh-16* and *ngn-1* reporter expression, using the same genomically-integrated transgenes, in *ngn-1* and *hlh-16* mutants, respectively. *hlh-16* reporter expression is not perturbed in *ngn-1(ns185)* mutants (5 experiments; n > 20 per time point per experiment; Figure 3E). However, *ngn-1* reporter levels are significantly reduced in *hlh-16(ns979)* mutants between 0-10 hours (n > 28 per time point; Figures 3A, S3B). Of note, *ngn-1* reporter levels recover to wild-type levels by 14 hours. Thus, *hlh-16* is required for *ngn-1* basal expression until 10 hours but is not required for the increase in *ngn-1* expression that follows. Consistent with these observations, we found that a high-copy *ngn-1p::ngn-1* cDNA transgene can restore Y-to-PDA transformation to *hlh-16* mutants (Figure 2P), supporting the notion that *hlh-16* functions to establish *ngn-1* basal expression levels.

To assess auto-regulation, we examined *ngn-1* and *hlh-16* reporter expression in *ngn-1* and *hlh-16* mutants, respectively. While *ngn-1* expression is not perturbed, *hlh-16* expression is significantly reduced in *hlh-16* mutants (6 experiments; n > 15 per time point per experiment; Figure 3E), suggesting that HLH-16 is required for its own expression.

### *egl-5/Hox* functions upstream of *ngn-1* and *hlh-16*

Previous studies showed that *egl-5/Hox* is required for Y-to PDA transformation. In *egl-5* mutants, a PDA neuron is not generated, and Y epithelial characteristics are constitutively maintained (Jarriault et al., 2008). To uncover possible interactions among *egl-5, hlh-16*, and *ngn-1, we* examined expression of the *ngn-1* and *hlh-16* reporters in two different *egl-5* mutants, *egl-5(n486)* and *egl-5(n945). We* found that both *egl-5* alleles strongly reduce *ngn-1* reporter expression at all time points, and induction of *ngn-1* expression between 10-14 hours is entirely blocked (n > 27 per time point, Figures 3A, S3C). Intriguingly, we found that expression of an *egl-5p::egl-5(exons1-3)::gfp* genomically-integrated reporter transgene (Ferreira et al., 1999) is dynamic over the imaging period. Between 0-12 hours, reporter levels do not vary extensively. However, after the 12-hour time point, there is a large increase in *egl-5* expression levels (4 experiments; n > 4 per time point per experiment; Figure S4B). This increase correlates with *ngn-1* reporter expression, raising the possibility that a rise in *egl-5* expression drives increased *ngn-1* expression. We believe that the onset of increased *egl-5* reporter expression is likely slightly delayed compared to the *ngn-1* reporter strain because of different growth rates of the strains. Our combined results strongly support the notion that EGL-5/HOX functions upstream of *ngn-1/Ngn* for both basal and induced expression, and suggest that these may be independent functions.

*egl-5* mutants also exhibit defects in *hlh-16* reporter expression. While initial levels of *hlh-16* appear normal, they steadily and significantly decline over the 14-hour imaging period (6 experiments; n > 17 per time point per experiment; Figure 3E). These observations suggest several conclusions. First, EGL-5 is not required for the initial expression of *hlh-16*. Second, EGL-5 is required for maintaining *hlh-16* gene expression before and during initiation of Y-to-PDA transformation. Third, the effects of *egl-5* on *ngn-1* are partly independent of *hlh-16*, as the levels of *hlh-16* at the start of the experiment appear normal, but *ngn-1* levels are dramatically reduced even at this time point. This last observation is consistent with our observations that *ngn-1* expression levels track the expression levels of *egl-5*, but not of *hlh-16*, between the 10-14 hour time points (Figures S4A,B).

### Temporal switching of SEM-4 function from a repressor to an activator of *ngn-1* expression

The *sem-4* gene encodes a SALL-family transcription factor previously reported to promote Y-to-PDA transformation, and we identified three alleles of the gene in our screen (Table S1). Like *egl-5* mutants, the Y cell in *sem-4* mutant animals retains epithelial properties and fails to transform into PDA (Jarriault et al., 2008). We therefore examined the expression of *ngn-1, hlh-16*, and *egl-5* reporter transgenes in *sem-4* mutants. Unexpectedly, and unlike in *egl-5* mutants, we found that in two different *sem-4* mutants (*ns1004, n1971)*, *ngn-1* reporter levels are induced to nearly twice the level of wild-type animals during the 0-10 hour observation period. This increased expression is subsequently retained, so that *ngn-1* expression levels at the 14-hour time point are comparable, or perhaps slightly higher than wild-type levels (n > 26 animals per time point; Figures 3A, S3D). These observations suggest that at early time points, SEM-4 normally represses *ngn-1* expression, but this inhibition becomes less pronounced as Y-cell morphology changes. Intriguingly, increased expression of *ngn-1* alone is not sufficient to drive PDA formation, as PDA formation is blocked even though *ngn-1* expression levels are high.

To understand the surprising effect of sem-4 loss on *ngn-1* expression, we tested whether *egl-5* and/or *hlh-16* are required for sem-4-dependent inhibition of *ngn-1* by generating CRISPR-mediated mutations in the *sem-4* gene in *egl-5(n945)* and *hlh-16(ns979)* mutant genetic backgrounds. We found that animals harboring loss-of-function mutations in both *egl-5* and *sem-4* have greatly reduced *ngn-1* reporter expression levels throughout the imaging period, comparable to *egl-5* single mutants alone (n > 24 per time point; Figure 3B). In *sem-4; hlh-16* double mutants, *ngn-1* reporter levels are slightly reduced compared to wild-type levels, as in *hlh-16* single mutants, during the 0-10 hour imaging period. However, *ngn-1* induction, which takes place normally in *hlh-16* single mutants, now fails to occur (n > 24 per time point; Figure 3C). Thus, from 0 to 10 hours, *sem-4* normally inhibits *ngn-1*, and the increased expression of *ngn-1* in *sem-4* mutants requires functional *egl-5* and *hlh-16* genes. At later times points, however, *sem-4* promotes *ngn-1* induction.

We also examined the effects of *sem-4* loss on *egl-5* and *hlh-16* reporter expression. Paradoxically, unlike *ngn-1, we* find that expression of both reporters is reduced (but not eliminated) at all time points (5 experiments, n > 15 per time point per experiment; n > 22 per time point; Figures 3E,S4B). Thus, *sem-4* promotes *egl-5* and *hlh-16* expression at all time points.

### LIN-12/NOTCH determines Y-to-PDA transformation timing through *ngn-1/Ngn* and *hlh-* 16/Olig-dependent and independent mechanisms

Our results strongly suggest that a temporal switch in Y-cell state, manifested by changes in the functions of the HLH-16 and SEM-4 transcription factors, correlates with initiation of Y-to-PDA transformation. We sought, therefore, to identify regulators of this cell-state change. A previous study reported that *lin-12(n676n930)* homozygous mutants, carrying a temperature-sensitive partial-loss-of-function mutation in the *lin-12/Notch* gene, fail to generate a Y cell at 25C (Jarriault et al., 2008). This allele was originally generated by reverting the *lin-12(n676)* gain-of-function allele to generate a *lin-12* allele containing the original gain-of-function mutation and a new loss-of-function mutation (Greenwald et al., 1983; Sundaram and Greenwald, 1993). We confirmed this result, showing that while a few newly-hatched L1 animals in this mutant at 25C express *hlh-16* or *ngn-1* reporters in a rectal cell with the morphology of the Y cell (Figure 4C), most do not (n > 70, Figure 4E). Curiously, in some animals, while a Y cell could not be detected, a PDA neuron expressing the *hlh-16* or *ngn-1* reporters was apparent. We wondered whether this precocious formation of PDA (>20 hours early) might reflect residual *lin-12/Notch* gain-of-function activity of this strain. We therefore examined newly-hatched animals at 15C, the permissive temperature for the loss-of-function defects, and found that almost all contain a precocious PDA neuron, and rare animals possess a normal Y cell (n > 80, Figures 4D,F). These results suggest that at 15C, *lin-12(n676n930)* animals have a fully-penetrant gain of *lin-12* function in Y. This is consistent with previous work showing that at 15C, this allele has some gain-of-function activity in other contexts (Sundaram and Greenwald, 1993).

**Figure 4:**
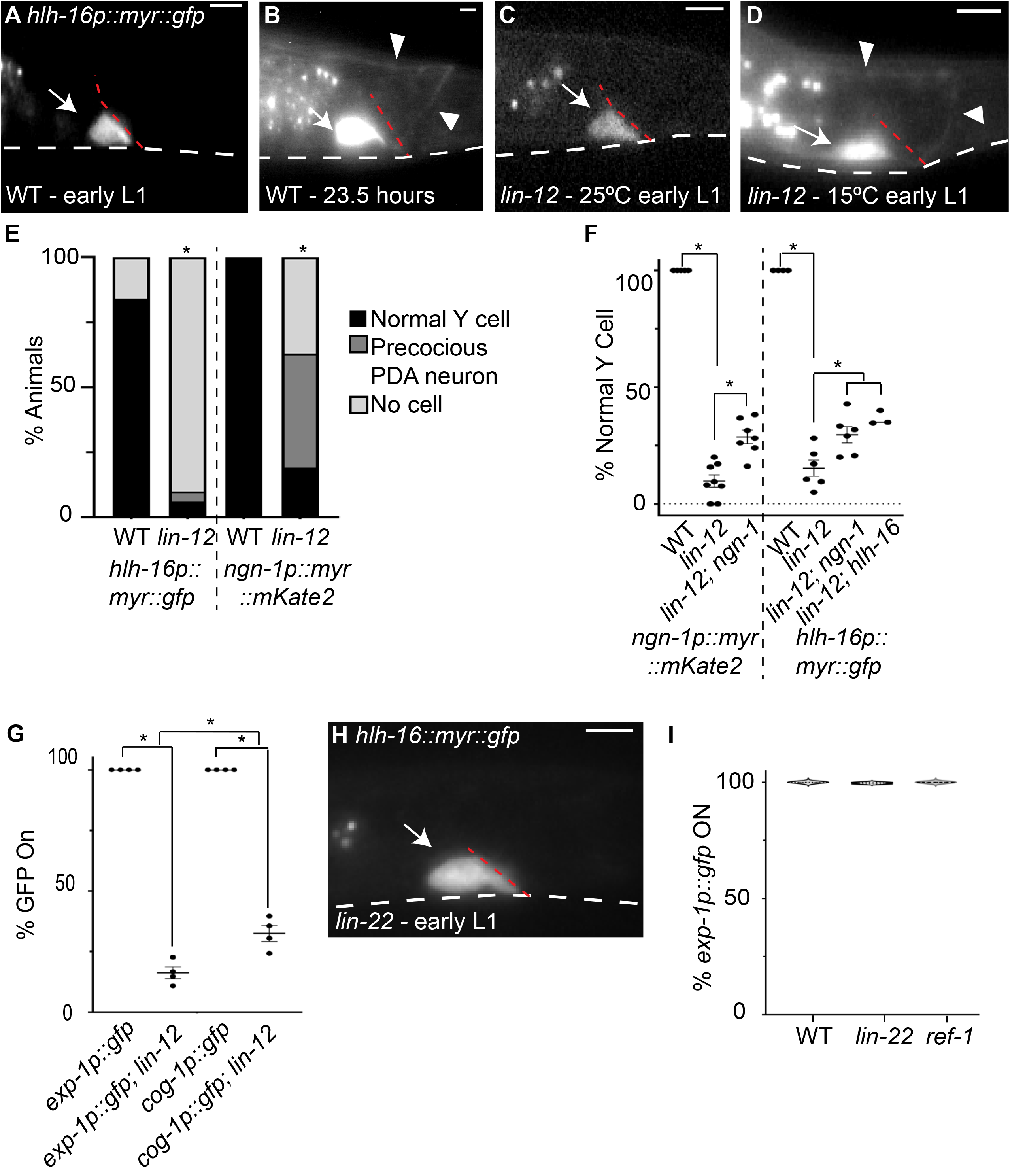
*lin-12/Notch* promotes Y-to-PDA transformation. (A) Wild-type (WT) Y cell in early L1 at 15C. (B) WT Y cell, 23.5 hrs after release from L1 arrest at 15C. (C) Early *lin-12(n676n930)* mutant L1 at 25C either has normal Y-cell morphology (shown) or does not express *nsIs943* [*hlh-16p::myr::gfp*]. (D) Early *lin-12(n676n930)* mutant L1 at 15C displays precocious PDA. (E) Quantification of Y/PDA in early L1 animals at 25C using *nsIs943* and *nsIs913* [*ngn-1p::myr::mKate2*] in WT and *lin-12(n676n930)* mutant animals. Fisher’s Exact Test, significance at p < 0.01. n > 70 animals. (F) Quantification of presence of a precocious PDA in early L1 animals at 15C using *nsIs913* and *nsIs943* reporter in WT, *lin-12(n676n930), lin-12(n676n930); ngn-1(ns988), lin-12(n676n930); ngn-1(ns185)*, and *lin-12(n676n930); hlh-16(ns196)* mutants. For *nsIs943*, percentages were quantified out of animals with fluorescent expression. One-way ANOVA with Tukey’s post-hoc test. Significance at p < 0.05. n > 50 animals for all genotypes. (G) Quantification of *nsIs130* [*exp-1p::gfp*] and *syIs63* [*cog-1p::gfp*] expression in WT and *lin-12(n676n930)* L4 animals raised at 15C. ANOVA with Tukey’s post-hoc test. Significance at p < 0.01. n > 100 animals for all genotypes. (H) Early *lin-22(n372)* mutant L1 at 20C with normal Y cell. (I) *nsIs131* [*exp-1p::gfp*] expression in WT and *lin-22(n372)* and *ref-1(mu220)* mutant L4 animals raised at 20C. One-way ANOVA. n > 350 animals. Scale bars, 5 μm. Dashed white lines, ventral surface of animal. Dashed red lines, rectal slit. Arrows, cell body. Arrowheads, axon.

To examine whether these precociously-formed PDA neurons are properly differentiated, we examined expression of *exp-1p::gfp* and *cog-1/Nkx6-3 promoter::gfp*, which is expressed slightly earlier (Zuryn et al., 2014), in *lin-12(n676n930)* animals grown at 15C. We found that only 18% of L4 animals express *exp-1p::gfp* and only 33% of L4 animals express *cog-1p::gfp* (n > 68; Figure 4G). Therefore, the Y cell in most *lin-12(n676n930)* animals executes stages I and II of Y-to-PDA transformation precociously but fails to reach stage III.

Is *ngn-1* required for the precocious formation of PDA? To address this question, we generated *lin-12(n676n930); ngn-1(ns988)* and *lin-12(n676n930); ngn-1(ns185)* double mutants and assayed *ngn-1* and *hlh-16* reporter expression respectively. Loss of *ngn-1* can suppress precocious PDA formation, suggesting that *lin-12/Notch* acts in part through ngn-1/neurogenin (n > 79; Figure 4F). Supporting this notion, a previous study reported that *lin-12/Notch* promotes *hlh-16* expression in other *C. elegans* motonerons (Bertrand et al., 2011), raising the possibility that NOTCH acts through *hlh-16* to affect *ngn-1* activity in Y/PDA. Indeed, loss of *hlh-16* can also suppress precocious PDA formation in *lin-12(n676n930)* mutants (n > 50; Figure 4F). Importantly, however, a substantial fraction of *lin-12(n676n930)* mutants, also carrying *ngn-1* or *hlh-16* mutations, still exhibit precocious PDAs, suggesting that a pathway independent of *ngn-1* must also exist.

In spinal-cord radial glia, NOTCH inhibits *Olig2* and *Ngn2* gene expression through HES-family transcriptional repressors (Sagner et al., 2018; Hatakeyama et al., 2004; Ishibashi et al., 1995; Ohtsuka, 1999). Our findings suggest that the opposite mechanism is at play during Y-to-PDA transformation, as *lin-12/Notch* promotes *hlh-16* and *ngn-1* activities. To understand this difference in more detail, we tested whether *lin-22*, encoding the only *C. elegans* protein containing all HES-protein domains (Wrischnik and Kenyon, 1997; Alper and Kenyon, 2001; Neves and Priess, 2005) or *ref-1*, a distantly related protein (Alper and Kenyon, 2001; Neves and Priess, 2005; Chou et al., 2015), are required for Y-to-PDA transformation. We found that *lin-22(n372)* and *ref-1(mu220)* mutants do not display precocious PDA formation (Figure 4H), and properly express the *exp-1p::gfp* reporter in PDA neurons of L4 animals (n > 350; Figure 4I). Thus, *lin-22* and *ref-1* are not required for Y-to-PDA transformation, consistent with a novel mechanism by which NOTCH regulates this epithelial-to-neuronal transition. Such a mechanism may share features with that employed in pancreatic islet development, where NOTCH promotes NGN3 expression (Cras-Méneur et al., 2009; Shih et al., 2012). The *C. elegans* Y/PDA may be a good system to study this elusive interaction.

Previous characterization of the *lin-12(n676)* strain argues that this allele increases wildtype LIN-12/NOTCH function (Greenwald et al., 1983; Sundaram and Greenwald, 1993). Taken together, our studies reveal that *lin-12/Notch* activity levels determine when Y-to-PDA transformation takes place. Low levels of NOTCH result in no transformation, which can be viewed as an infinite delay in transformation onset; wild-type levels result in transformation initiation at 10 hours post L1-arrest; and excess *lin-12/Notch* expression drives precocious PDA formation.

### Cytoskeletal organizers UNC-119, UNC-44/ANK and UNC-33/CRMP function downstream of HLH-16 and NGN-1

Our findings that *exp-1/Gabrb3* reporter expression turns on after the onset of PDA axon extension, and that *lin-12(gf)* results in formation of a precocious axon-bearing PDA neuron defective in *exp-1/Gabrb3* expression, prompted us to investigate how the transition from stage II to stage III of Y-to-PDA transformation takes place. To address this, we went back to the mutants collected in our screen. As with other mutants we found, *ns200* animals exhibit a defect in *exp-1p::gfp* expression, but generate a normal Y cell in L1 animals (Figure 5A). Unlike *ns185* and *ns196* mutants, *ns200* mutants correctly silence the *egl-5p::gfp* reporter over development, suggesting that *ns200* mutants may be defective in transitioning from stage II to stage III of Y-to-PDA transformation (Figure 5A). *ns200* mutants also exhibit pronounced locomotion defects. SNP mapping uncovered linkage of the causal lesion to chromosome III, and whole-genome sequencing revealed a G-to-T mutation in the *unc-119* gene (Figure 5C). This mutation is predicted to cause a G79Stop mutation, truncating all isoforms of the UNC-119 protein. Introducing the G79Stop lesion into the genomic *unc-119* locus of wild-type animals via CRISPR (allele *ns913)*, reproduces the *exp-1/Gabrb3* expression defect of *ns200* mutants, although the penetrance of the defect is lower (Figure 5A), suggesting that *unc-119(ns200)* mutants also carry enhancer mutations in the background. A transgene containing 1 kb of *unc-119* upstream regulatory sequences, fused to the *unc-119* cDNA from the related nematode *C. briggsae*, restores both *exp-1/Gabrb3* expression and wild-type locomotion to transgenic animals (Figure 5B). Furthermore, an *ngn-1p::unc-119* cDNA transgene fully restores *exp-1/Gabrb3* expression, but only weakly rescues the locomotion defects of *unc-119* mutants (Figure 5B), suggesting that *unc-119* acts cell autonomously in Y/PDA.

**Figure 5:**
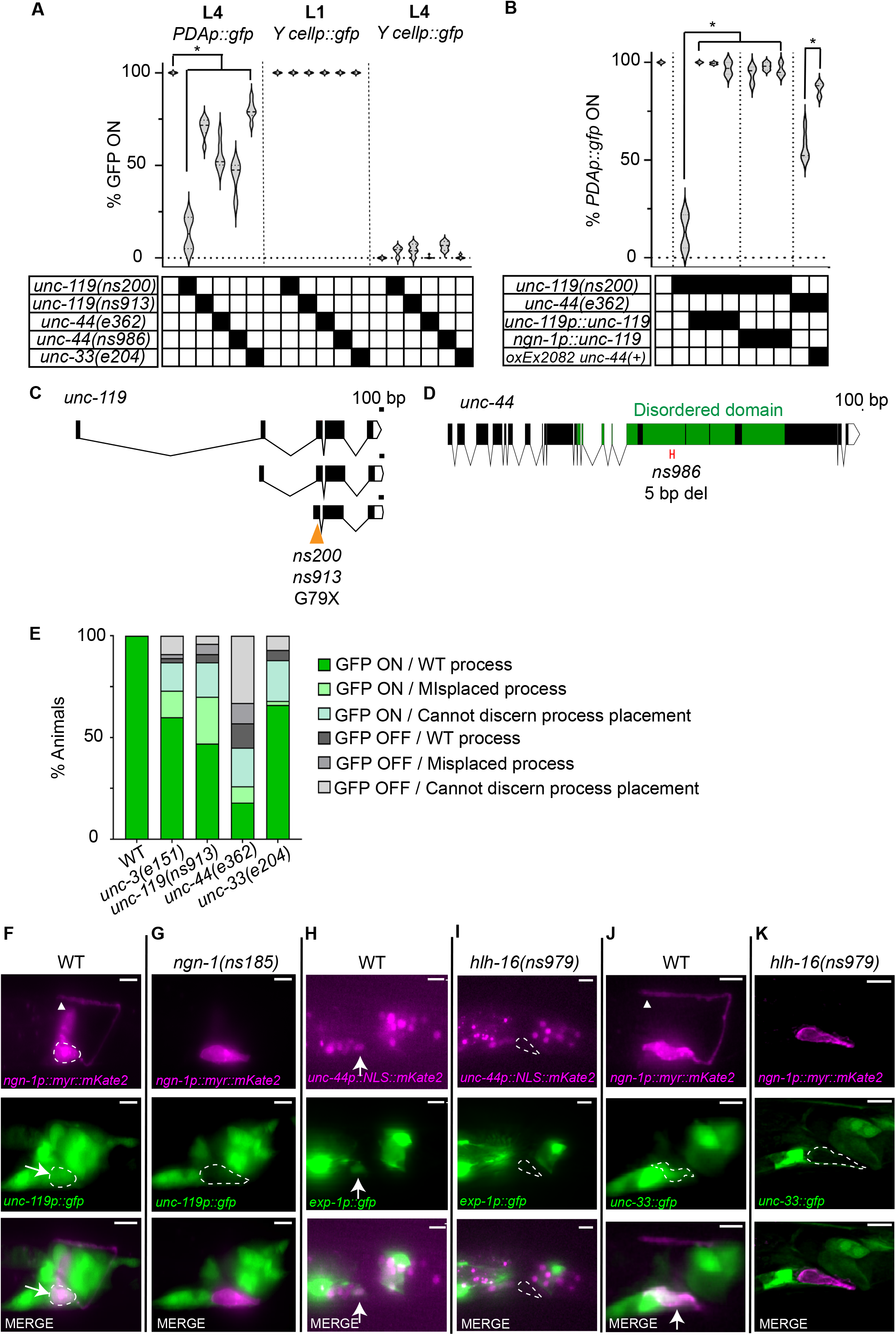
Cytoskeleton organizers UNC-119, UNC-44, and UNC-33 promote *exp-1p::gfp* expression in the PDA neuron. See also Figure S5; Table S2. (A) *unc-119, unc-44*, and *unc-33* mutants are defective in PDA maturation. Two-way ANOVA with Dunnett’s post-hoc test. Significance at p < 0.01. n > 100 animals. (B) Similar to Figure 2P. *PDAp::gfp, nsIs131*[*exp-1p::gfp*]. A one-way ANOVA with Dunnett’s post-hoc test was performed for comparisons with *unc-119* mutants, and a t-test was performed for comparison with *unc-44*. Significance at p < 0.01. n > 68 animals. (C-D) Amino acid changes are indicated. X, stop codon. (D) Only the long neuronal isoform of *unc-44* is shown. *ns986* has a 5 bp deletion nucleotides 15,987 to 15,992, leading to an early stop codon. (E) Defects in *PDAp::gfp* expression are independent of axon guidance defects. n > 100. (F-K) Expression of *unc-119, unc-44*, and *unc-33* in wild type and indicated mutants. Arrows, cell body. Arrowhead, axon tip.

When we introduced the *ngn-1* and *exp-1/Gabrb3* reporters together into *unc-119(ns913)* mutants, we found that, unlike the other mutants we studied, most animals extend a PDA axon. In one third of the animals, the process is abnormal, and in about 13% of the animals, *exp-1/Gabrb3* expression is not detected (n > 100; Figures 5E,S5). Our results, therefore, demonstrate that the *unc-119* gene is required for the transition from stage II to stage III.

UNC-119 is a highly conserved protein that acts as a cytoskeleton organizer in neurons (Maduro et al., 2000; Knobel et al., 2001; Manning et al., 2004; Materi and Pilgrim, 2005; He et al., 2020). A recent study reported that *unc-119* functions together with *unc-44/Ankyrin* and *unc-33/Crmp* to regulate dendrite polarization, likely through control of actin and microtubule organization, respectively (Hedgecock et al., 1985; Otsuka et al., 1995; Otsuka et al., 2002; Boontrakulpoontawee and Otsuka, 2002; Tsuboi et al., 2005; Zhou et al., 2008; Maniar et al., 2011; Norris et al., 2014; He et al., 2020; LaBella et al., 2020; Chen et al., 2021). Consistent with this, we found that animals carrying mutations in either *unc-44/Ank* or *unc-33/Crmp* exhibit PDA differentiation defects similar to those seen in *unc-119* mutants (Figure 5A). A loss-of-function mutation, *ns986*, specifically affecting the neuronal isoform of *unc-44*, also displays PDA differentiation defects, and a full-length extrachromosomal *unc-44* gene (*oxEx2082;* LaBella et al., 2020) can rescue *unc-44(e362)* mutant PDA defects (Figures 5A,B,D). Thus, surprisingly, cytoskeletal regulators are important for proper gene expression in stage III of Y-to-PDA transformation.

How might such regulation occur? Although most *unc-119, unc-44*, or *unc-33* mutants have defective PDA axons (Figure 5E), in all three mutants, a population of animals with fully formed, properly placed PDA axons are found that fail to express the *exp-1/Gabrb3* reporter (Figures 5E,S5). This observation suggests that the functions of *unc-119/unc-33/unc-44* in axon extension and *exp-1/Gabrb3* expression are at least partly independent.

To understand when *unc-119, unc-33*, and *unc-44* expression begins during Y-to-PDA transformation, we examined respective reporter constructs. Reporter expression for all three genes is first visible in PDA after the axon begins to grow (Figure 5F,H,J). Importantly, in *hlh-16* or *ngn-1* mutants, these reporters are not expressed at all (Figure 5G,I,K). Thus, *unc-119, unc-33*, and *unc-44* are likely direct or indirect targets of *ngn-1. We* found that mutations in the COE-type transcription factor gene *unc-3* have axon extension and *exp-1/Gabrb3* expression defects highly reminiscent of those seen in *unc-119, unc-33*, and *unc-44* mutants (Figure 5E). This is consistent with previous observations (Richard et al., 2011), and suggests the possibility that NGN-1 might indirectly affect the expression of cytoskeleton-organizing genes through UNC-3.

## DISCUSSION

### A model for the control of Y-to-PDA transformation

The data presented here are consistent with the model for Y-to-PDA transformation depicted in Figure 6. We propose that NGN-1/NGN plays a pivotal role in the decision of Y to undergo differentiation. Early on, LIN-12/NOTCH activates HLH-16/OLIG. HLH-16/OLIG levels are then maintained by auto-activation and EGL-5/HOX, which is, in turn, activated by SEM-4/SALL. NGN-1/NGN is positively regulated by HLH-16/OLIG but also inhibited by SEM-4/SALL, keeping its expression at a basal level. At later time points, NGN-1/NGN is released from control by SEM-4/SALL and HLH-16/OLIG, and its expression levels are boosted through increased EGL-5/HOX expression, requiring SEM-4/SALL, at least in part. Following the increase in NGN-1/NGN, expression of the cytoskeleton regulators UNC-119, UNC-33/CRMP, and UNC-44/ANK is induced, directly or indirectly by NGN-1. UNC-119, UNC-33/CRMP, and UNC-44/ANK are required for PDA axon extension, but also for transcription of the *exp-1/Gabrb3* GABA receptor gene, and perhaps other downstream PDA maturation genes.

**Figure 6:**
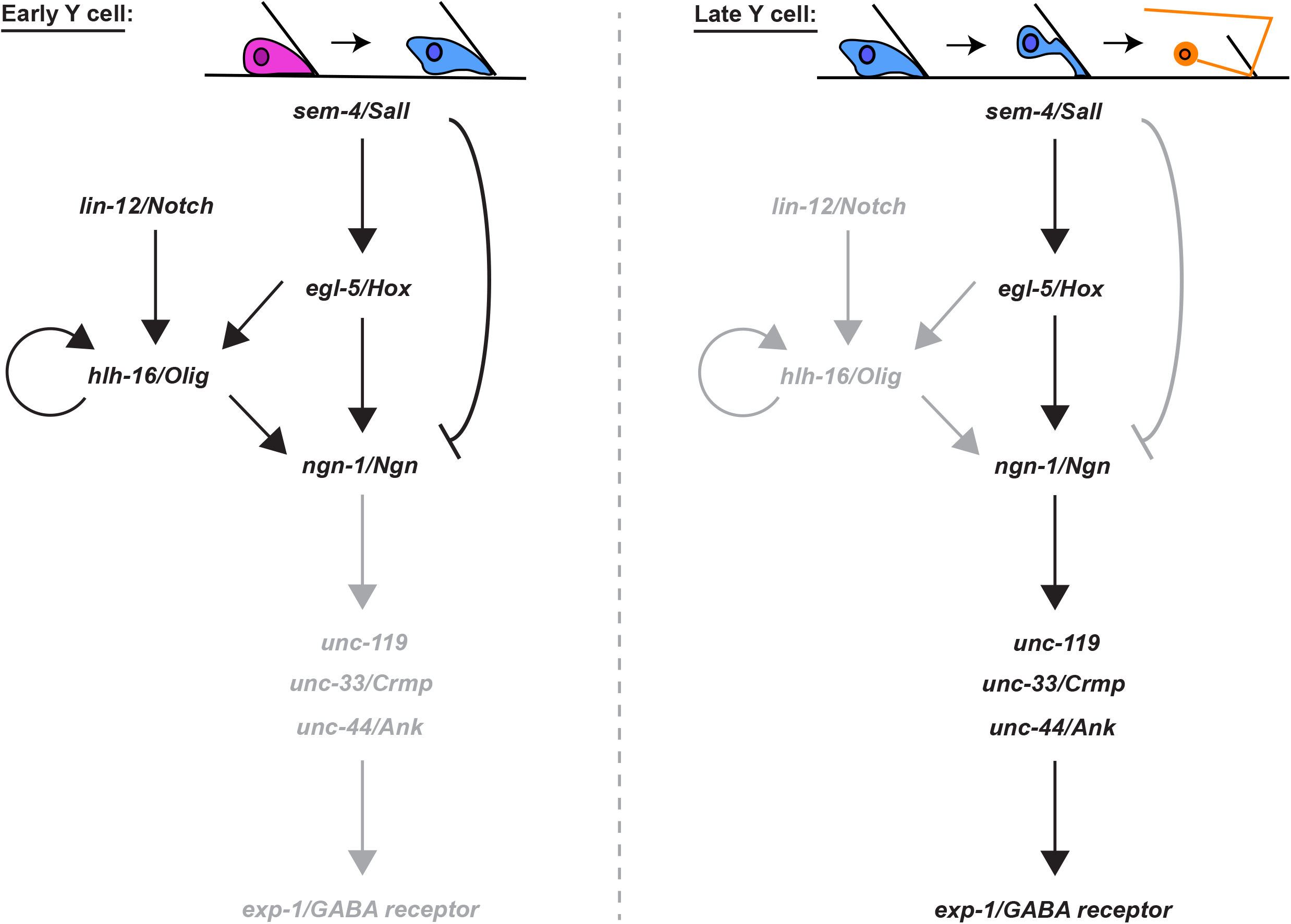
A model for Y-to-PDA transformation. Black and gray letters indicate active or inactive genes/interactions, respectively.

Our model posits that NGN-1/NGN is key for PDA formation in wild-type animals. However, two results suggest that there are likely additional means by which PDA can be produced: precocious PDA formation in *lin-12/Notch* mutants is only partially blocked by loss of *ngn-1*, and PDA formation is blocked in *sem-4/Sall* mutants despite high levels of *ngn-1* expression. The latter observation could also be explained if NGN-1 becomes inactivated in *sem-4* mutants. In mammals, some bHLH proteins are rapidly inactivated by phosphorylation when they reach peak expression (Quan et al., 2016).

Our model also suggests that SEM-4/SALL has opposing activities as a transcriptional activator and repressor. Previous studies suggest that SEM-4 is part of the NODE complex, including a number of transcription factors that together activate *egl-5* in the Y cell (Kagias et al., 2012). In other settings, however, SALL proteins have been suggested to be repressors (Ott et al., 2001; Barembaum and Bronner-Fraser, 2004; Parrish et al., 2004; Sweetman and Münsterberg, 2006). Whether both activities co-exist in the Y cell, or whether there is a switch is not clear.

Finally, we point out that the events depicted in Figure 6 occur in the absence of cell division, suggesting that similar cell-division-independent roles for homologous proteins in other animals are likely.

### A conserved module for epithelial-to-motoneuron/motoneuron-like fate transitions

The similarities between Y-to-PDA transformation, spinal-cord motoneuron formation, and pancreatic islet cell generation are striking. All three events begin with a precursor cell situated within a tube-lining epithelium. Why tubular structures are appropriate sites for progenitor cells is not clear. While pancreatic ducts and the spinal canal could represent milieus that contain signaling molecules for stem-cell differentiation (Huang et al., 2010; Lehtinen et al., 2011; Chang et al., 2012; Feliciano et al., 2014), this is unlikely to be the case for the Y cell, where the rectal canal likely transports only intestinal waste. Alternatively, tube-lining epithelial cells may be under specific mechanical stresses that facilitate stem-cell division or maintenance (Park et al., 2017; Bogdanova et al., 2018; Guerrero et al., 2019; Legøy et al., 2020).

All three events also rely on *Neurogenin-family* members to promote differentiation (Ma et al., 1999; Fode et al., 1998; Mizuguchi et al., 2001; Lee et al., 2005; Zhou et al., 2007; Rubio-Cabezas et al., 2011; Gradwohl et al., 2000; Bankaitis et al., 2018), and Olig-family bHLH proteins are also important. The latter have been directly implicated in vertebrate motoneuron generation and in PDA formation (Takebayashi et al., 2000; Novitch et al., 2001; Mizuguchi et al., 2001; Takebayashi et al., 2002; Lu et al., 2002; Lee et al., 2005). In the pancreas, expression of *bHLHb4*, an Olig-family bHLH transcription factor, is reported in cells bordering pancreatic ducts (Bramblett et al., 2002); however, no functional studies of the protein in the context of islet formation have been published.

*Notch* also plays a commanding role in all three differentiation events, although our studies suggest that the effects are not the same in all contexts. In the spinal cord and pancreas, NOTCH, acting through HES transcription factors, inhibits differentiation of stem cells. In *C. elegans*, NOTCH activity in Y promotes PDA formation independently of the LIN-22/HES and REF-1/HES-related protein. A group of nematode-specific proteins distantly related to HES proteins have been described in the REF-1 family to function with NOTCH during *C. elegans* embryogenesis (Alper and Kenyon, 2001; Neves and Priess, 2005; Chou et al., 2015). It is possible that one of these other family members is a relevant NOTCH target, and that instead of inhibiting gene expression, they promote transcription, explaining the difference with the mammalian settings. Whatever the mechanism, it is likely conserved, as similar NOTCH-dependent, HES-independent induction of NGN3 is seen in the pancreatic islets (Cras-Méneur et al., 2009; Shih et al., 2012). Thus, future investigation of Y-to-PDA transformation could provide additional insight into mammalian pancreatic differentiation.

SALL proteins related to SEM-4 participate in early neural differentiation in the brain, are required for neuronal migration in chick neural crest cells, suppress chick neurogenin, and are expressed in the mouse spinal-cord ventricular zone (Ott et al., 2001; Barembaum and Bronner-Fraser, 2004; Parrish et al., 2004; Sweetman and Münsterberg, 2006), suggesting that these proteins may indeed be relevant for spinal-cord motoneuron and/or pancreatic islet development. However, their activities in these contexts have not been assessed. Similarly, roles for HOX proteins in neural and pancreatic development have been demonstrated (Aigner and Gage, 2005; Li et al., 2017; Tan et al., 2016; Catela et al., 2016; Larsen et al., 2015; Shi, 2010), but specific interactions with the Neurogenin pathway have not been extensively studied.

### Cytoskeletal organizers as regulators of gene expression

Our finding that genes controlling cytoskeleton organization are required for transcriptional maturation of the PDA motoneuron is surprising, as it is not immediately clear how these cytoplasmic proteins can affect nuclear events. Recent studies demonstrate that ankyrin proteins, related to *C. elegans* UNC-44, are required for proper pancreatic islet function (Healy et al., 2010), and ankyrins have been studied extensively for their roles in neuronal cytoskeleton organization and protein filtering into neuronal processes (Nakada et al., 2003; Jenkins et al., 2015; Sobotzik et al., 2009; Dubreuil and Yu, 1994; Jegla et al., 2016; LaBella et al., 2020). Could these roles be tied to transcriptional control? One possibility is that cellular polarity establishment, which requires cytoskeletal components for localizing proteins that initiate maturation, activates transcription factors that promote expression of terminal-differentiation genes. In neuronal contexts, the cytoskeleton could also monitor synapse maturation (LaBella et al., 2020), setting up a retrograde signal that influences nuclear transcription. Such retrograde signaling has been described in several contexts (Hendry et al., 1974; Feldman et al., 1999; Scheiffele et al., 2000; Asmus et al., 2000; Poo, 2001; E. Alger, 2002; Baudet et al., 2008). Given the facility with which we identified mutants defective in Y-to-PDA transformation, it is possible that the mechanisms at play could be revealed through additional forward genetic approaches in *C. elegans*.

## Acknowledgments

We would like to thank Scott Emmons, Iva Greenwald, Maxwell Heiman, Oliver Hobert, and Erik Jorgensen for strains. Some strains were provided by the CGC, funded by NIH Office of Research Infrastructure Programs (P40 OD010440). Images were acquired at the Bio-Imaging Resource Center, RRID:SCR_017791, and Whole Genome Sequencing was performed at the Genomics Resource Center at The Rockefeller University. We would like to acknowledge the Electron Microscope Imaging Facility of CUNY Advanced Science Research Center for instrument use, and scientific and technical assistance. Electron microscopy was performed at the New York Structural Biology Center, supported by grants from the Simons Foundation (349247), NYSTAR, and NIH (GM103310). This work was funded by a Kavli pilot grant to A.R., a fellowship from the NYSCF to M.T., an NSF predoctoral fellowship to A.R., and NIH grant R35 NS105094 to S.S.

## Author Contributions

Conceptualization, A.R., M.T., S.S.; Methodology, A.R. and S.S.; Validation, A.R. and M.T.; Investigation, A.R. and M.T.; Resources, S.S.; Writing – Original Draft, A.R.; Writing – Review & Editing, A.R., M.T., and S.S.; Visualization, A.R.; Electron Microscopy, Y.L.; Supervision, S.S.; Project Administration, S.S.; Funding Acquisition, S.S.

## SUPPLEMENTAL FIGURE LEGENDS

**Figure S1:**
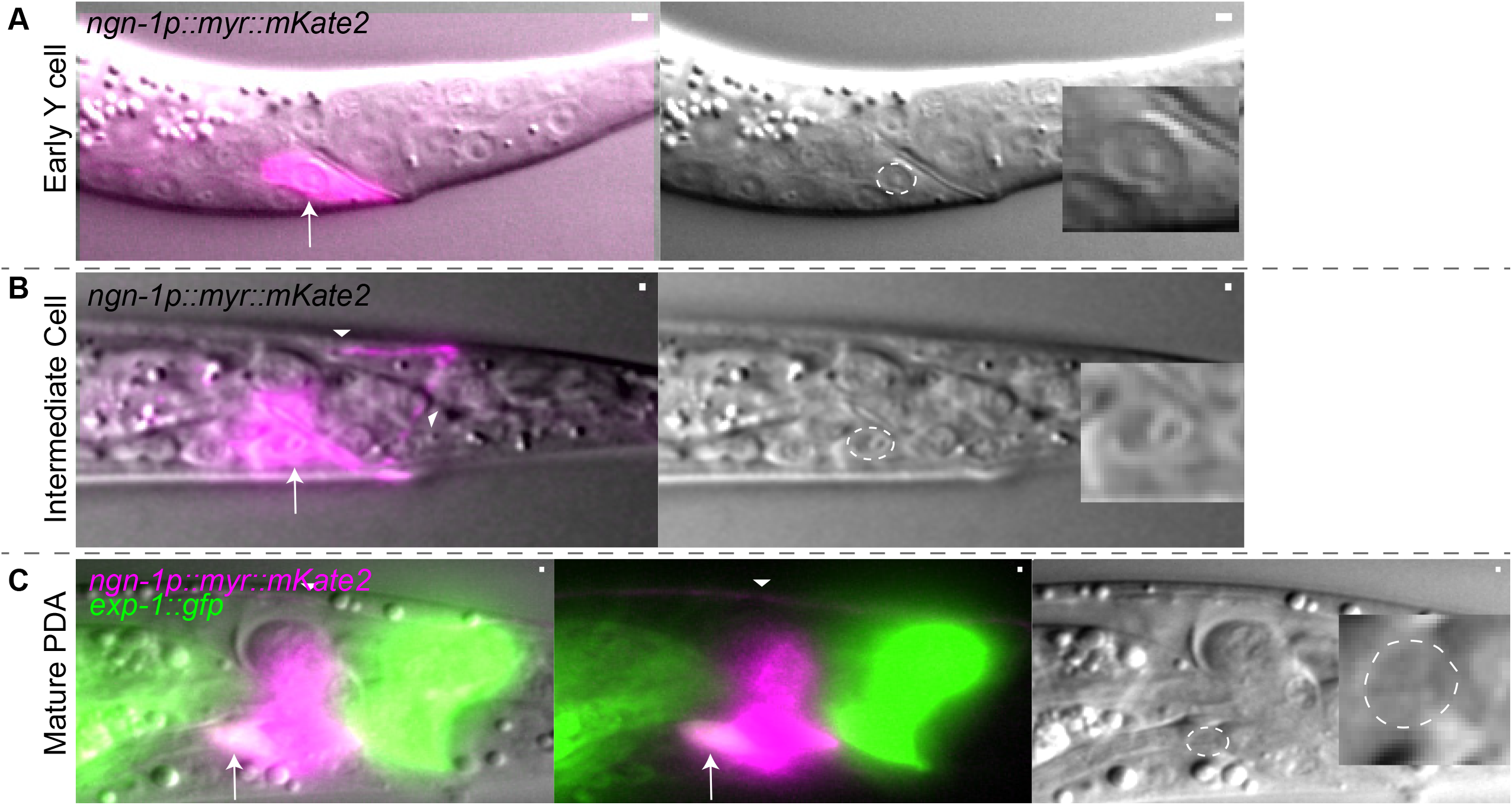
PDA adopts a neuron-like speckled nuclear morphology after axon outgrowth is complete. Related to Figure 1. (A) Early Y cell has a “fried egg” epithelial-like nucleolar morphology. (B) Y/PDA with a process still has a “fried egg” nucleolar morphology. (C) Mature PDA (with *exp-1p::gfp* expression) adopts a neuron-like speckled nucleolar morphology. Arrows, cell body. Arrowheads, cellular process. Dashed circle, outline of cell body. Scale bars, 5 μm.

**Figure S2:**
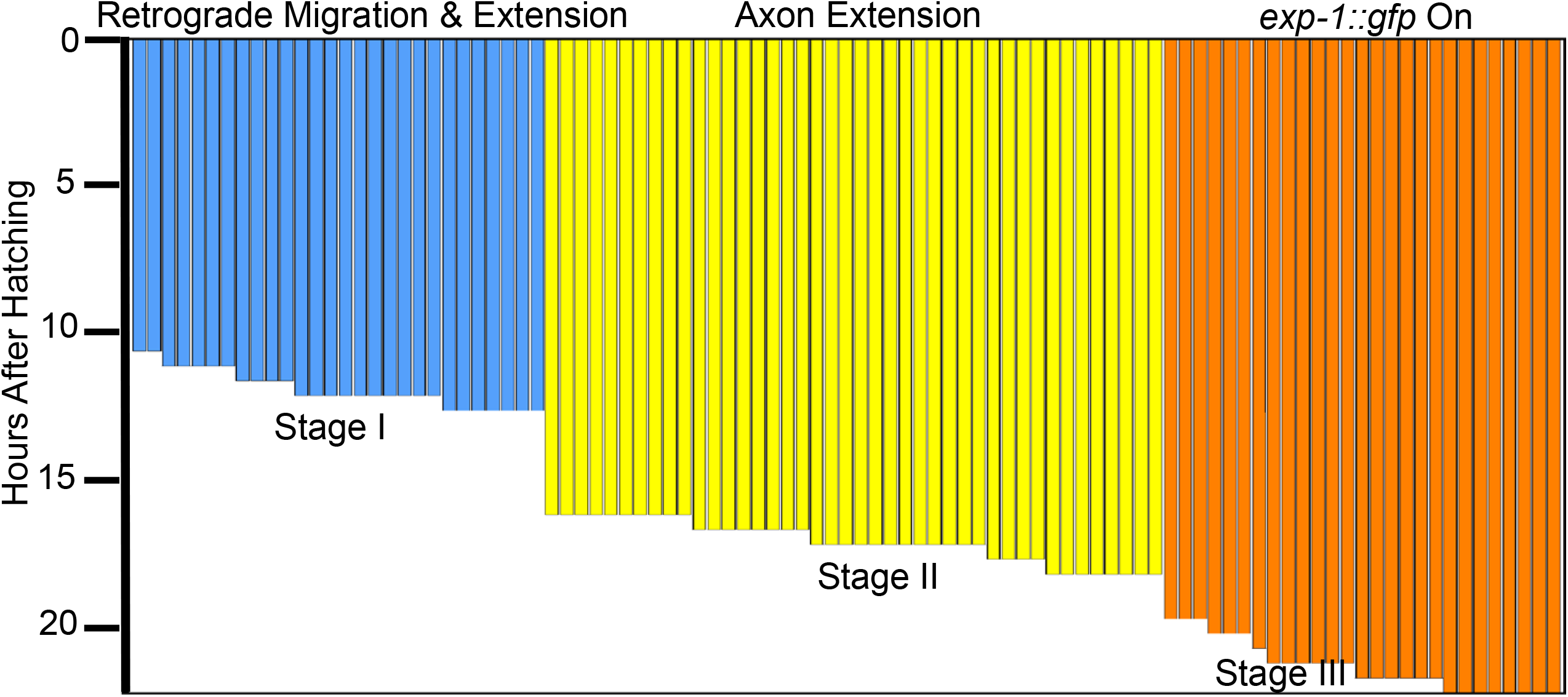
Starvation does not affect the timing or nature of Y-to-PDA transformation events. Related to Figure 1. Each horizontal bar represents Y-to-PDA stage of a single animal. Stage I: n = 28, mean = 11.77 ± 0.63 hrs, Stage II: n = 42, mean = 16.9 ± .71 hrs, Stage III: n = 27, mean = 21.11 ±0.86 hrs.

**Figure S3:**
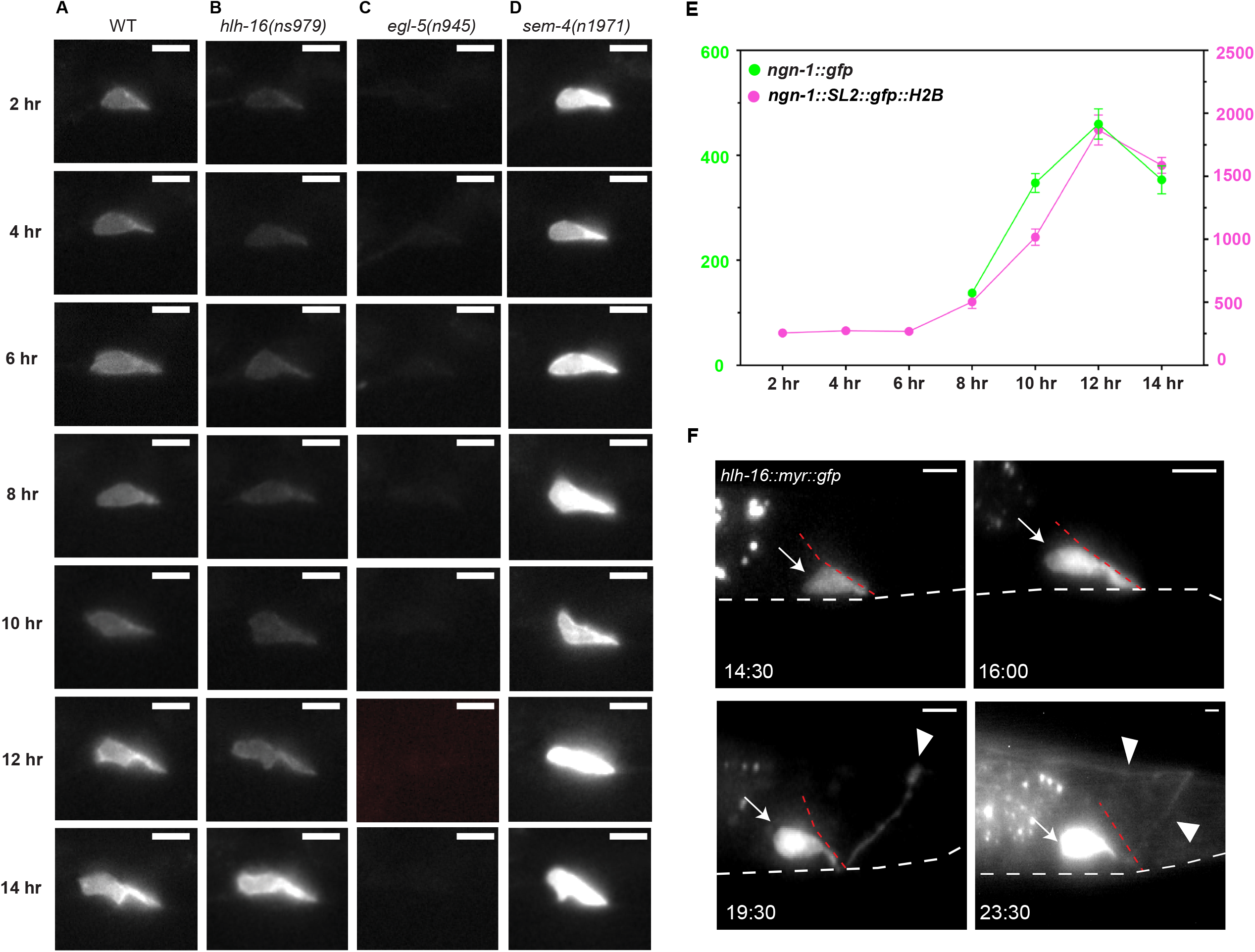
Following Y-to-PDA transformation using *ngn-1::myr::mKate2* and *hlh-16p::myr::gfp::hlh-16 3’utr*. Related to Figure 3. Time stamps indicate times after release from L1 arrest. Scale bars, 5 μm. (A-D) Representative images of Y cells in indicated mutant backgrounds used for quantification of Figure 3A. (E) Time course of *ngn-1(syb5802)* endogenous translational gfp fusion and *ngn-1(syb5813)* endogenous transcriptional histone fused gfp. *ngn-1(syb5802)* expression is not detectable before the 8-hour time point and is detectable in 6/13 animals at the 8-hour timepoint. n > 11 animals per time point for each genotype. (F) Following Y-to-PDA transformation using *hlh-16p::myr::gfp::hlh-16 3’utr*. (Top left) Y cell before transformation begins. (Top right) Retrograde cell-body migration and formation of thick process that remains attached to rectal epithelium. (Bottom left) Axon extension. (Bottom right) PDA process extension is complete. Note that *hlh-16* expression becomes fainter at this time point. Arrow, cell body, arrowhead, axon. White dashed line, ventral surface of animal. Dashed red line, rectal slit.

**Figure S4:**
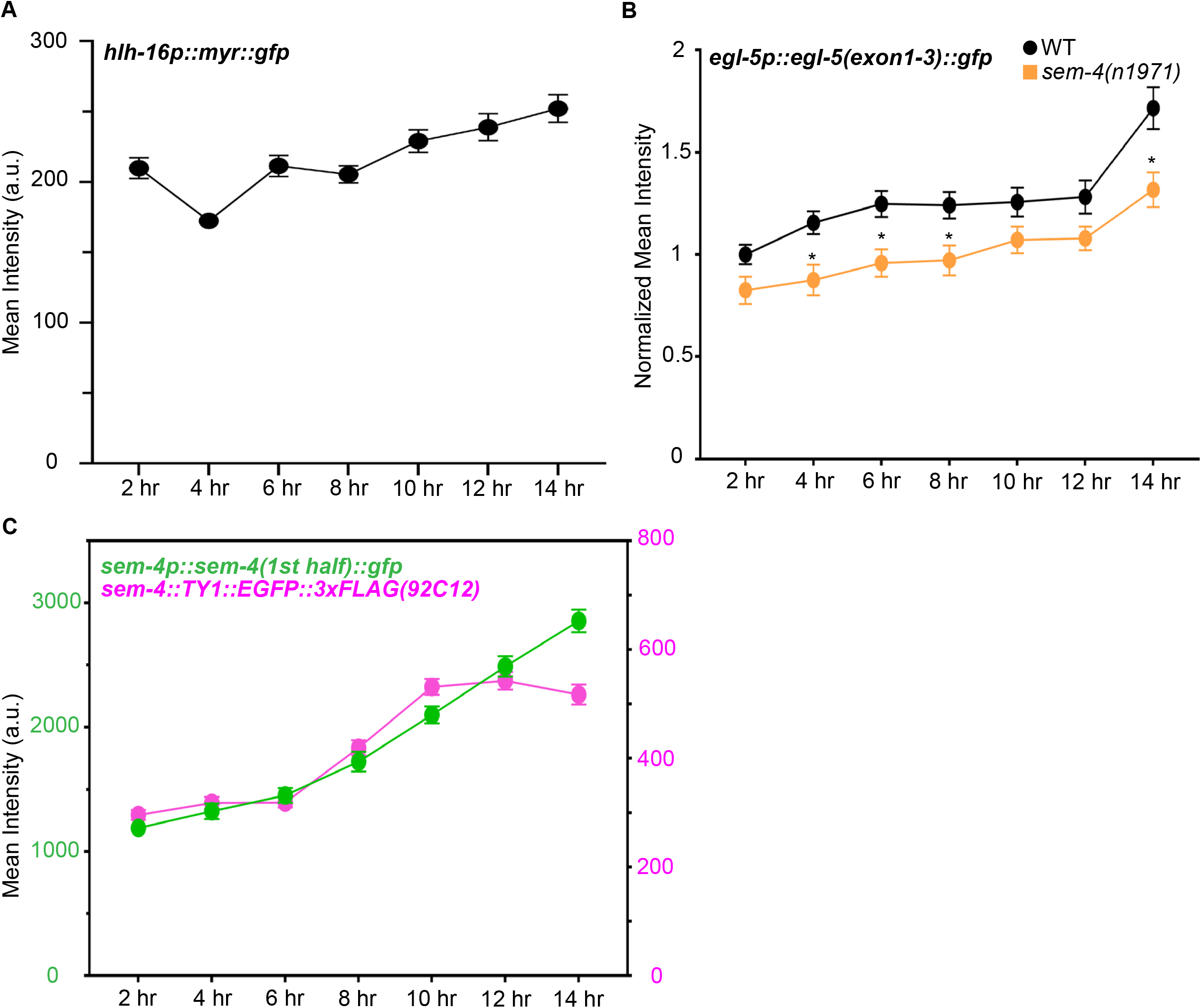
*hlh-16, egl-5*, and *sem-4* reporter expression in early stages of Y-to-PDA transformation. Related to Figure 3. (A) GFP fluorescence intensity in wild-type animals harboring an *nsIs943* [*hlh-16p::myr::gfp*] reporter transgene. Mean grey value intensity at the center slice of an 80 pixel x 80 pixel region containing the Y cell. n > 50 animals for each time point. (B) GFP fluorescence intensity in animals expressing a *bxIs7* [*egl-5p::egl-5(exon1-3)::gfp*] reporter in the indicated genotypes. Mean grey value intensity in the center slice in an ROI drawn around the cell and normalized to the average values for the corresponding wild type at the 2 hr timepoint. n > 23 animals for each genotype per time point. t-tests were performed at each time point. Significance at p < 0.01. (C) GFP fluorescence intensity in wild-type animals expressing *kuIs34* [*sem-4p::sem-4(5’ half)::gfp*] and *wgIs57* [sem-4::TY1::EGFP::3xFLAG(92C12)] reporters. Mean grey value intensity in the center slice in and ROI drawn around the cell body. Error bars, SEM. n > 24 animals for each genotype per time point.

**Figure S5:**
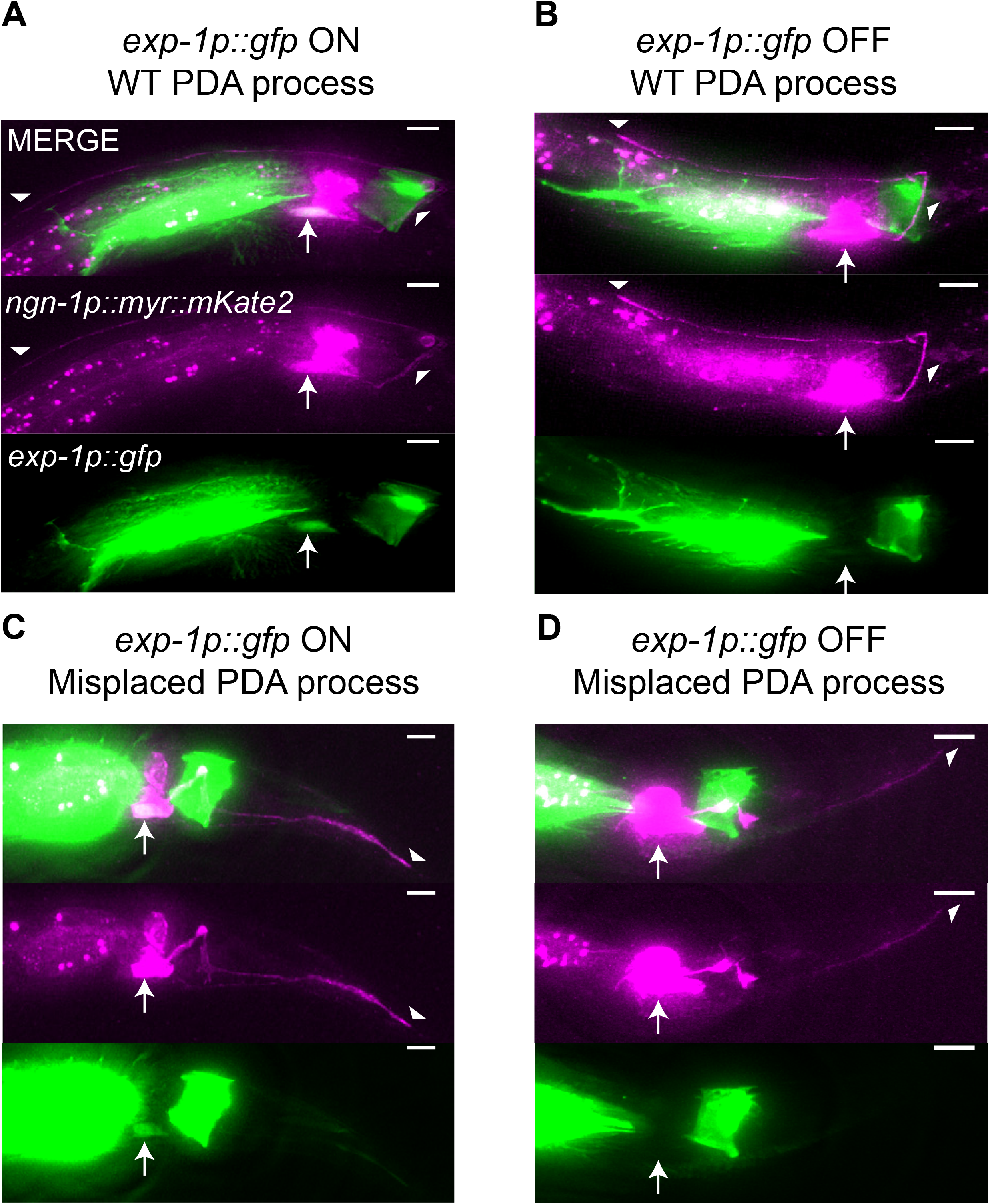
Defects observed in mutants of the cytoskeleton organizing genes *unc-119, unc-44*, and *unc-33*. Related to Figure 5. (A) A wild-type PDA mutant in an *unc-44(e362)* mutant. The PDA process is properly placed and *exp-1p::gfp* expression is on. (B) *unc-44(e362)* mutant; PDA process is properly placed, but *exp-1p::gfp* is not expressed. (C) *unc-44(e362)* mutant, PDA process is improperly placed, but *exp-1p::gfp* is expressed. (D) PDA process is improperly placed, and *exp-1p::gfp* is not expressed. Arrows, cell body. Arrowheads, cellular process. Scale bars, 5 μm.

## STAR METHODS

### RESOURCE AVAILABILITY

#### Lead Contact

Further information and requests for resources and reagents should be directed to the lead contact, Shai Shaham.

#### Materials Availability

All unique strains and reagents generated in this study are available from the lead contact.

#### Data and Code Availability

Data and statistical analyses are available in Supplementary Table 2.

### EXPERIMENTAL MODEL AND SUBJECT DETAILS

#### Mutants and Reporters

Experiments were conducted at 20C unless otherwise indicated. *C. elegans* were cultivated using standard methods (Brenner, 1974). The wild-type parent for most strains used in this study is the *C. elegans* Bristol strain N2. The relevant mutations used in this study are: LG I: *hlh-16(ns196), hlh-16(ns989), sem-4(n1971)*; LG II: *ref-1(mu220)*; LG III: *egl-5(n945), egl-5(n486), lin-12(n676n930), unc-119(ns200), unc-119(ns913);* LG IV: *unc-44(e362), unc-44(ns986), unc-44(syb28170), ngn-1(ns186), ngn-1(syb2810), ngn-1(ns988), ngn-1(syb5802), ngn-1(syb5813), lin-22(n372)*. Transgenes used in these studies were *nsIs131* [3.5 *kb exp-1p::gfp::unc-54 3’utr*] III, *bxIs7* [*egl-5p::egl-5(exon1-3)::gfp*], *syIs63* [*cog-1::GFP*], *otIs45* [*unc-119::gfp*]; *otls118* [*unc-33::gfp*], *nsIs913* [*4 kb ngn-1p::myr::mKate2::unc-54 3’utr*] IV, *nsIs914* [*4 kb ngn-1p::myr::mKate2::unc-54 3’utr*], *nsIs942* [*4 kb ngn-1p::myr::mKate2::unc-54 3’utr; 4 kb ngn-1::ajm-1::yfp::unc-54 3’utr*], *nsIs943* [480 bp *hlh-16p::myr::GFP::1.5 kb hlh-16 3’utr*], *wgIs57* [*sem-4::TY1::EGFP::3xFLAG(92C12) + unc-119(+)*], *kuIs34* [*sem-4::GFP*], *oxEx2082* [*unc-44 long isoform rescue*]. Further information provided in Table S3.

### METHOD DETAILS

#### Electron Microscopy

L1 animals with short Y/PDA processes as in Figure 1F were selected for electron microscopy based on *ngn-1p::myr::mKate2* fluorescence signals in the tail. Animals were prepared and sectioned using standard methods (Lundquist et al., 2001). Serial images were acquired by using a Titan Themis 200 kV transmission electron microscope with Cs Image Corrector. Image processing and analysis were performed using NIH ImageJ and IMOD software (Schorb et al., 2019).

#### Plasmid construction

Plasmids were constructed using Gibson cloning (New England Biolabs) into the applicable backbone plasmid.

#### Mutant analysis and scoring

An EMS screen was performed in the *nsIs131* background, seeking F2 L4 animals lacking *exp-1p::gfp* expression in PDA (Jorgensen and Mango, 2002). Isolates were transferred to individual plates. Penetrance for each mutant was scored for several generations. Once causative mutations were identified in the alleles, *bxIs7* [*egl-5p::egl-5(exon1-3)::gfp*] was used to assess the formation of the Y cell in L1 animals and the absence of the Y cell in L4 animals.

#### SNP Mapping

For SNP mapping, mutants were crossed to CB4856 Hawaiian (HA) animals. F2 animals lacking *exp-1p::gfp* expression were isolated and assessed for the percent penetrance to confirm that it matches the penetrance from the parent strain. F2 recombinants were lysed and analyzed for three SNPs between N2 and HA backgrounds on each chromosome. If a chromosomal region showed linkage to N2 DNA, further SNPs were used to narrow down the chromosomal region of the mutation.

#### Whole genome sequencing

Animals were grown on *E. coli* strain OP50 until bacteria were depleted, harvested in M9, and resuspended in 1 M pH 7.5 TEN. SDS and proteinase K were added to a final concentration of 0.5% and 0.1 mg/ml respectively and 1 μl of β-mercaptoethanol was also added. The reaction was incubated overnight in a shaking thermocycler at 56C. Phenol/chloroform was added and phase-separated by spinning in a phase-lock tube. The aqueous phase was transferred to 200 proof EtOH. The resulting DNA clot was washed in 70% EtOH, dried, and resuspended in TEN. After rehydration, 0.3 μl of 100 mg/ml RNAse A was added and incubated at 37C for 2 hours. Phenol/chloroform extraction was performed again and DNA was rehydrated with EB buffer. The sample was then run on a nanodrop machine and on a 1% gel to confirm that there was no RNA contamination or DNA degradation, respectively. NextSeq High Output 1 x 75 sequencing was performed with Standard Illumina Sequencing primers for gDNA-seq application.

#### Germline Transformation and Rescue Experiments

Plasmid mixes containing the plasmid of interest, co-injection markers, and pBluescript were injected into both gonads of young adult hermaphrodites at a total of 100 ng/ul. Injected animals were singled onto NGM plates and allowed to grow for two generations. Transformed animals based on co-injection markers were picked onto single plates and screened for stable inheritance of the extrachromosomal array. Only lines from different P0 injected hermaphrodites were considered independent. Lines with stable inheritance of the extrachromosomal array were used to screen for rescues.

#### CRISPR Cas9 Genome Editing

Alleles of *ngn-1(ns988), hlh-16(ns979), unc-119(ns913)*, and *unc-44(ns986)* were generated using a co-CRISPR editing strategy (Farboud et al., 2019). *sem-4(ns1004-1008)* mutants were generated using a co-injection strategy (Dokshin et al., 2018). Guide crRNA, repair single-stranded DNA oligos, tracrRNA, and buffers were ordered from IDT. Guide crRNAs used are listed below.

*hlh-16:* 5’ CATTGTTACTAGCTTCCAATTGG 3’
*unc-119:* 5’ GCTCTTCCAGGAATCACTCAAGG 3’
*unc-44:* 5’ GTAGTTTCCATAACATGACGCGG 3’
*ngn-1:* 5’ TCGCCAGTTTGCAAATGATGCGG 3’
*sem-4:* 5’ TGCAAAGACAAATGGCGAGCCGG 3’

5’ ATTGAAAGTGTTTACAGAGAGG 3’

#### Timing of Y-to-PDA transformation events

To determine the timing of Y-to-PDA events, *nsIs913; nsIs131* gravid hermaphrodites were bleached and then incubated on a rotator in M9 overnight for L1 arrest synchronization. L1s were then plated onto agar plates containing NGM medium and *E. coli* OP50 bacteria. We imaged separate groups from this pool every 30 minutes between 6-22 hours after food introduction, and scored for retrograde cell body migration, process extension, and onset of *exp-1p::gfp* expression until all animals imaged completed all three events.

#### Quantification of fluorescence in the Y cell

To quantify the expression of various reporters before process extension, gravid hermaphrodites were bleached and then incubated on a rotator in M9 overnight for L1 arrest synchronization. L1s were then plated onto agar plates containing NGM medium and *E. coli* OP50 bacteria. Separate groups were imaged every 2 hours until 14 hours after release from L1 arrest. *nsIs913* [*ngn-1p::myr::mKate2*], *nsIs943* [*hlh-16p::myr::gfp::hlh-16 utr*], *wgIs57* [*sem-4::TY1::EGFP::3xFLAG*], and *kuIs34* [*sem-4::gfp*] animals were imaged on a Zeiss Imager M2 with an AxioCam r1.4 on a 63x/1.4 NA oil objective at 50, 150, 50, 30 ms exposure respectively. For *nsIs913* and *nsIs943*, an 80 pixels x 80 pixels ROI was drawn over the Y cell. The fluorescence intensity was quantified at the z-position at the center of the cell. For *wgIs57* and *kuIs34*, an ROI was drawn around the Y cell nucleus, and the fluorescence intensity was quantified at the z-position at the center of the cell body. *bxIs7* was imaged on an iSIM with a VisiTech iSIM head and a Leica DMi8 inverted microscope stand with a 100x/1.47 NA oil objective and an Orca sCMOS camera. An ROI was drawn over the entire Y cell, and the fluorescence intensity was quantified at the z-position at the center of the cell.

#### lin-12 temperature-sensitive experiments

*lin-12(n676n930)* mutant animals were raised at 15C. Gravid hermaphrodites were bleached and then incubated on a rotator in M9 at 15C or 20C overnight for L1 arrest synchronization. L1s were plated onto agar plates containing NGM and immediately prepared for imaging.

#### Cytoskeleton organizers mutant morphology

Gravid animals were bleached and then incubated on a rotator in M9 overnight for L1 arrest synchronization. L1s were then plated onto agar plates containing NGM medium and *E. coli* OP50 bacteria. 47 hours later, animals were prepared for imaging.

#### UNC-44 expression localization

A CRISPR insertion of *SL2::NLS::mKate2* before the stop codon in the long neural isoform of *unc-44* was obtained from SUNY Biotech (Fuzhou, China).

#### Imaging preparation

2% agarose with 6 mM NaN_3_ was used to make thin pads for mounting animals on microscope slides. Animals were picked into a 2 μl drop of M9 and a 0.17 mm coverslip was placed on top. *ngn-1p::myr::mKate2* animals for figure images were imaged on an inverted Olympus IX-70 microscope with a 60X silicone objective, a pco.edge sCMOS camera, and deconvolution using measured PSFs.

### QUANTIFICATION AND STATISTICAL ANALYSIS

GraphPad Prism 9.2.0 was used to perform statistical analysis on the data. Specific statistical details can be found in the figure legends. Error bars represent SEM. Statistical significance was determined using p < 0.05 or p < 0.01. Data and statistical results for each figure can be found in Supplementary Table 2.

