## Supplementary material for "A developmental pathway for epithelial-to-motoneuron transformation in *C. elegans*": Table S1

**Table S1:** **Results of a forward genetics screen for loss of *exp-1p::gfp* expression.** Related to Figure 2.

| Gene Name | Allele | Number of Isolated Alleles | Molecular Lesions |
| --- | --- | --- | --- |
| *hlh-16* | *ns196* | 1 | R50C, See Figure 3 |
| *mab-9* | *ns187*  *ns206* | 2 | G282D |
| *ngn-1* | *ns185*  *ns194* | 2 | Q17X, See Figure 3  Q29X |
| *sem-4* | *ns188*  *ns189*  *ns192* | 3 | S520G  S520G  G295X |
| *unc-119* | *ns200* | 1 | G79X, See Figure 3 |
| Unidentified | *ns184*  *ns186*  *ns193*  *ns195*  *ns197*  *ns199*  *ns201* | 7 |  |
